# Structure of the ciliary axoneme at nanometer resolution reconstructed by TYGRESS

**DOI:** 10.1101/363317

**Authors:** Kangkang Song, Zhiguo Shang, Xiaofeng Fu, Xiaochu Lou, Nikolaus Grigorieff, Daniela Nicastro

**Affiliations:** Departments of Cell Biology and Biophysics, University of Texas Southwestern Medical School, Dallas, TX 75390, USA.; Janelia Farm Research Campus, Howard Hughes Medical Institute, 19700 Helix Drive, Ashburn, VA 20147, USA.

## Abstract

The resolution of subtomogram averages calculated from cryo-electron tomograms (cryo-ET) of crowded cellular environments is often limited due to signal loss in, and misalignment of the subtomograms. In contrast, single-particle cryo-electron microcopy (SP-cryo-EM) routinely reaches near-atomic resolution of isolated complexes. We developed a novel hybrid-method called “<u>T</u>omograph<u>Y</u>-<u>G</u>uided 3D <u>RE</u>construction of <u>S</u>ubcellular <u>S</u>tructures” (TYGRESS) that combines cryo-ET with SP-cryo-EM to achieve close-to-nanometer resolution of complexes inside crowded environments. Using TYGRESS, we determined the native 3D structures of the intact ciliary axoneme with up to 12 Å resolution. These results reveal many structures and details that were not visible by cryo-ET. TYGRESS is generally applicable to cellular complexes that are amenable to subtomogram averaging, bringing us a step closer to (pseudo-)atomic models of cells.

**One Sentence Summary:** A hybrid cryo-electron microscopy method reveals subcellular structures at unprecedented resolution.

## Main Text

Due to recent hardware and image processing advances, single-particle (SP-) cryo-EM can produce 3D reconstructions of purified, native proteins and macromolecular complexes (from ~50 kDa to several MDa size) with near-atomic detail, i.e. with a resolution of 3 Å or better (Bai et al. 2015, Cheng 2015, Danev and Baumeister 2017). Even single-particle cryo-ET of relatively thin (<100 nm) samples containing isolated complexes (Bartesaghi et al. 2012), and of viruses with a high abundance of capsomers (Schur et al. 2016, Wan et al. 2017) has reached sub-nanometer resolution. However, a few hundred nanometer thick and complex cellular samples are not presently amenable to structural study by SP-cryo-EM. The reason is that for unambiguous particle picking and accurate particle alignment, SP-cryo-EM usually requires the particles to be purified, structurally relatively homogeneous and distributed in a thin monolayer that avoids superposition of particles in the cryo-EM projection images (Cheng et al. 2015).

Cryo-ET is unique in that it can 3D-reconstruct and visualize pleomorphic structures, such as intact cells and organelles *in situ.* In addition, repeating identical components in the reconstructed tomograms, such as axonemal repeats in cilia or chemoreceptors in bacterial membranes, can be resolved with molecular detail using subtomogram averaging that increased the signal-to-noise ratio and thus resolution of the reconstruction (Nicastro et al. 2006, Briggs 2013, Fu et al. 2014). However, the resolution of cellular cryo-ET and subtomogram averaging is ultimately limited by the need to balance several irreconcilable factors (table S1). For example, a higher electron dose improves the signal-to-noise ratio of the tilt images, increasing accuracy of image alignment and correction of the contrast transfer function (CTF), with positive effects on resolution. On the other hand, a higher electron dose also leads to more structural degradation by radiation damage, limiting useful high-resolution signal to the early exposures in a tomogram. In contrast to thin “single particle-type” samples, the signal in tomograms of cellular samples is degraded by: a) increased inelastic electron scattering due to the large thickness of cellular samples and the effective thickness increasing by 1/*cis*(*α*) with tilt angle *α*; b) incomplete sampling of the reconstruction in Fourier space beyond a given resolution (Crowther criterion: m = π ^∗^ D/d, with number of tilt images m, sample thickness D and resolution d) (Crowther et al. 1970); c) beam-induced motion and electrostatic charging affecting images of tilted samples more severely; and d) the initial fast, not fully correctable motion in exposures is reiterated with every new exposure in a tilt series (Brilot et al. 2012). Some efforts have been made to optimize these factors to achieve higher resolution with subtomogram averaging, such as “constrained single-particle cryo-electron tomography” that uses constrained projection-matching refinement procedures (Bartesaghi et al. 2012, Yu et al. 2013, Himes and Zhang 2017), and dose-symmetric tilt-schemes combined with exposure filtering to more efficiently use the high-resolution information from early exposure images that contain less radiation damage (Grant and Grigorieff 2015, Hagen et al. 2017). However, only averaging subtomograms of relatively thin and uncrowded samples produces nanometer (or better) resolution (Bartesaghi et al. 2012, Schur et al. 2016, Wan et al. 2017). Subcellular samples are usually more crowded and thicker than these samples. It is therefore difficult to process their micrographs and novel strategies are required to push the resolution of *in situ* imaging to close the gap to high-resolution structure determination methods.

## TYGRESS is a hybrid method for *in situ* structural studies

Here we describe a novel hybrid-method called “TomographY-Guided 3D REconstruction of Subcellular Structures” (TYGRESS) [Fig. 1, and supporting online materials (SOM) text] to resolve structures *in situ* in crowded cellular environments with higher resolution than was previously possible using cryo-ET and subtomogram averaging. TYGRESS is essentially a single-particle reconstruction from untilted high-dose (HD) images recorded with an electron dose typical for SP-cryo-EM (30-60 e/A^2^), which is 10-60 times higher than the electron dose used for individual low-dose (LD) images of a tilt series (Fig. 1A). The HD images therefore contain optimally preserved high-resolution signal by using single exposures of untilted specimen. The SP-cryo-EM reconstruction of protein assemblies in their complex cellular context is made possible by providing additional 3D information for particle picking and initial particle alignment from the cryo-ET reconstruction and subsequent subtomogram averaging that is performed on the same specimen area where a HD image is recorded (Fig. 1, B and C).

**Figure 1.**
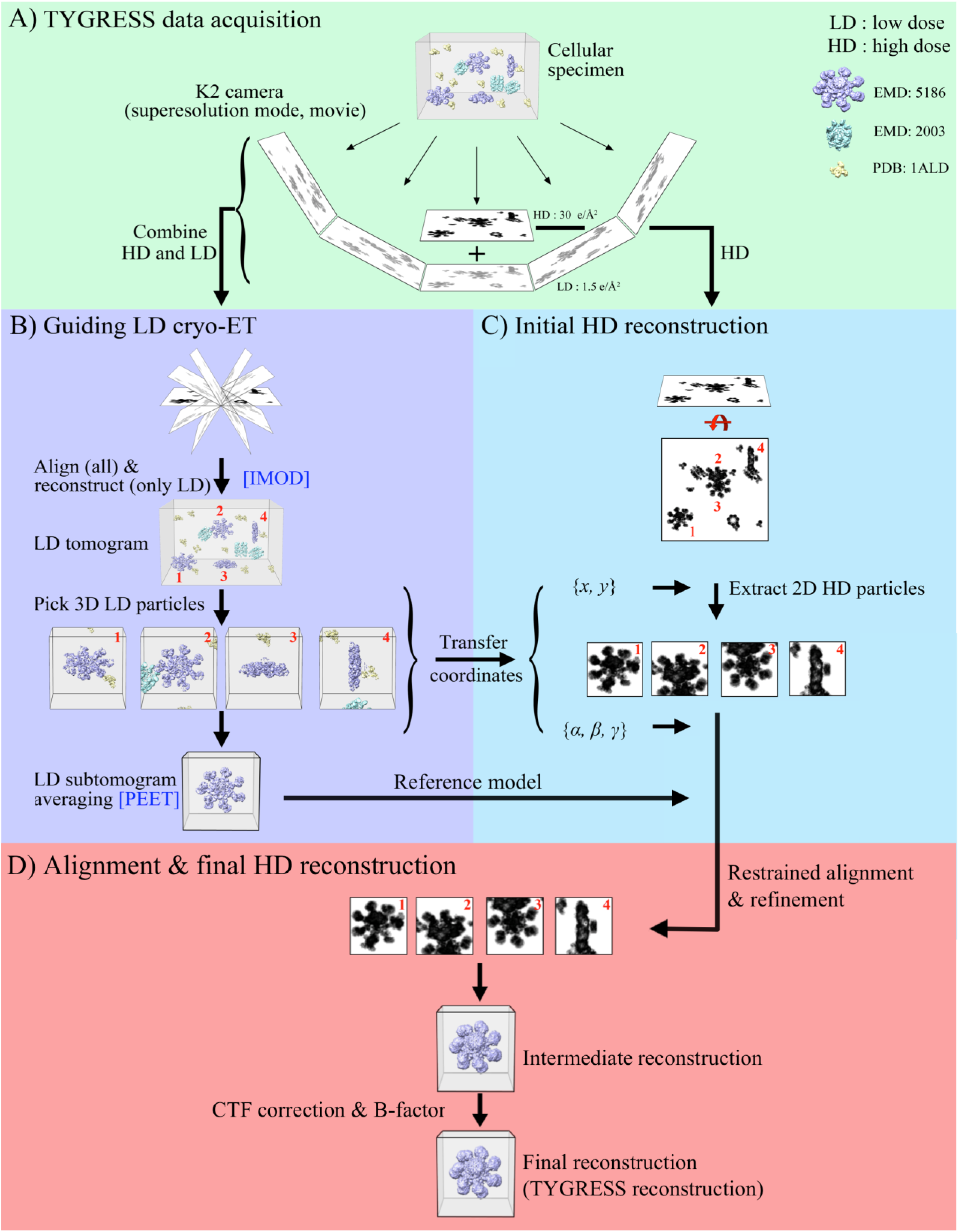
Overview of the TYGRESS workflow. To simulate a ‘cellular specimen’, cryo-EM structures of apoptosomes (EMD: 5186, purple), GroEL (EMD: 2003, cyan), and aldolase (EMD: 1ALD, yellow), were randomly placed in a 3D volume. A. A conventional single-particle image (HD image, ~30 e/Å^2^) is recorded, followed by a tomographic tilt series (~1.5 e/Å^2^ per LD image) of the same sample.
B. Both the LD images and the HD image are aligned. After alignment, only the LD images are used to reconstruct a tomogram, and particles of interest are picked and subjected to subtomogram averaging.
C. Particles of interest are picked in the HD image using the coordinates {*x*, *y*}_SP_ and orientation (*α*, *β*, *γ*}_SP_ derived from the tomogram in ***B***.
D. The particles picked from the HD image are aligned using their corresponding orientations and locations of the subtomograms. A CTF-corrected single-particle reconstruction is calculated from the particles in the HD image and used to further refine particle orientations and locations. Finally, a negative B-factor is applied to visualize high-resolution details.

Briefly, the TYGRESS workflow includes: (1) for each sample area two datasets are acquired, a HD image at 0° followed by a typical cryo-ET tilt-series (Fig. 1A); (2) the cryo-tomograms are reconstructed followed by subtomogram averaging of the particles of interest (Fig. 1B); (3) the information of the 3D particle position in the tomogram is projected onto the 2D HD image for particle picking (Fig. 1C and Fig. S1), and the subtomogram angles for each particle are applied to the respective particles extracted from the HD image for initial particle alignment (Fig. 1C); finally (4) the particles extracted from the HD images are further processed, which includes constrained single particle-type alignment, refinement for sub-averaged particles, and CTF- and B-factor correction to generate the final reconstruction (Fig. 1D and Fig. S2), whereas the cryo-ET data themselves are not included in the final TYGRESS average. There are multiple advantages of TYGRESS over cryo-ET/subtomogram averaging, including previously published strategies for resolution improvement, which in sum should lead to a considerable resolution improvement: a) the higher electron dose of the HD image significantly improves the signal-to-noise ratio and thus the reliability of SP-cryo-EM refinement strategies and CTF correction; b) HD images are recorded with an electron does of ~30 e/Å^2^ (at 300 kV) and therefore are affected far less by radiation damage than images used in regular cryo-ET/subtomogram averages, which suffer accumulated doses of up to ~100 e/ Å^2^), and c) the image quality of HD images is not degraded by sample tilting and multiple exposures.

## Validation of TYGRESS using the ciliary axoneme as model system

For validation we applied TYGRESS to an important and highly complex cell organelle that has been well-studied by cryo-ET and subtomogram averaging, the intact ciliary axoneme from the multiciliated protist *Tetrahymena thermophila.* The axoneme is the evolutionarily conserved microtubule core of cilia and flagella with a canonical array of nine outer doublet microtubules (DMTs) surrounding two central single microtubules that form the scaffold for more than 400 different associated proteins (Fig. S3) (Pazour et al. 2005). Notably, each DMT is composed of 96-nm repeat units, which can be treated as particles for subtomogram averaging. In the more than 30 published cryo-ET studies of the >200 nm thick ciliary axonemes the achieved resolution ranged from ~3-4 nm resolution (FSC=0.5) (Nicastro et al. 2006, Lin et al. 2014, Lin et al. 2014, Lin and Nicastro 2018), with the best resolution of 2.5 nm (FSC=0.5) being reached recently using Volta-Phase-Plate and K2 data (Fu et al. 2018). These studies provided important insights into the functional organization of axonemes, the mechanisms of dyneins and cilia motility, and ciliary dysfunction in human diseases (Nicastro et al. 2006, Lin et al. 2014, Lin and Nicastro 2018). However, the resolution of cryo-ET is still insufficient to characterize molecular interactions and regulatory pathways in greater details.

By applying TYGRESS to intact *Tetrahymena* axonemes, we obtained an averaged 3D structure of the 96-nm axonemal repeat *in situ* with considerably improved resolution of up to 12 Å (FSC=0.143) (Figs. 2 to 4, Figs. S4 to S6, and Movie S1). Many previously unseen molecular complexes and structural details *in situ* are revealed, such as the 96-nm ruler and its interactions with other axonemal rulers (Fig. 3, A to E, and Fig. S4), individual N-DRC components (Fig. 3F and Fig. S4) and newly identified microtubule inner proteins (MIPs) (Fig. 4, and Fig. S5 and S6). This allows a better understanding of the molecular mechanisms of ciliary assembly and the roles played by individual axonemal proteins in normal ciliary function.

**Figure 2.**
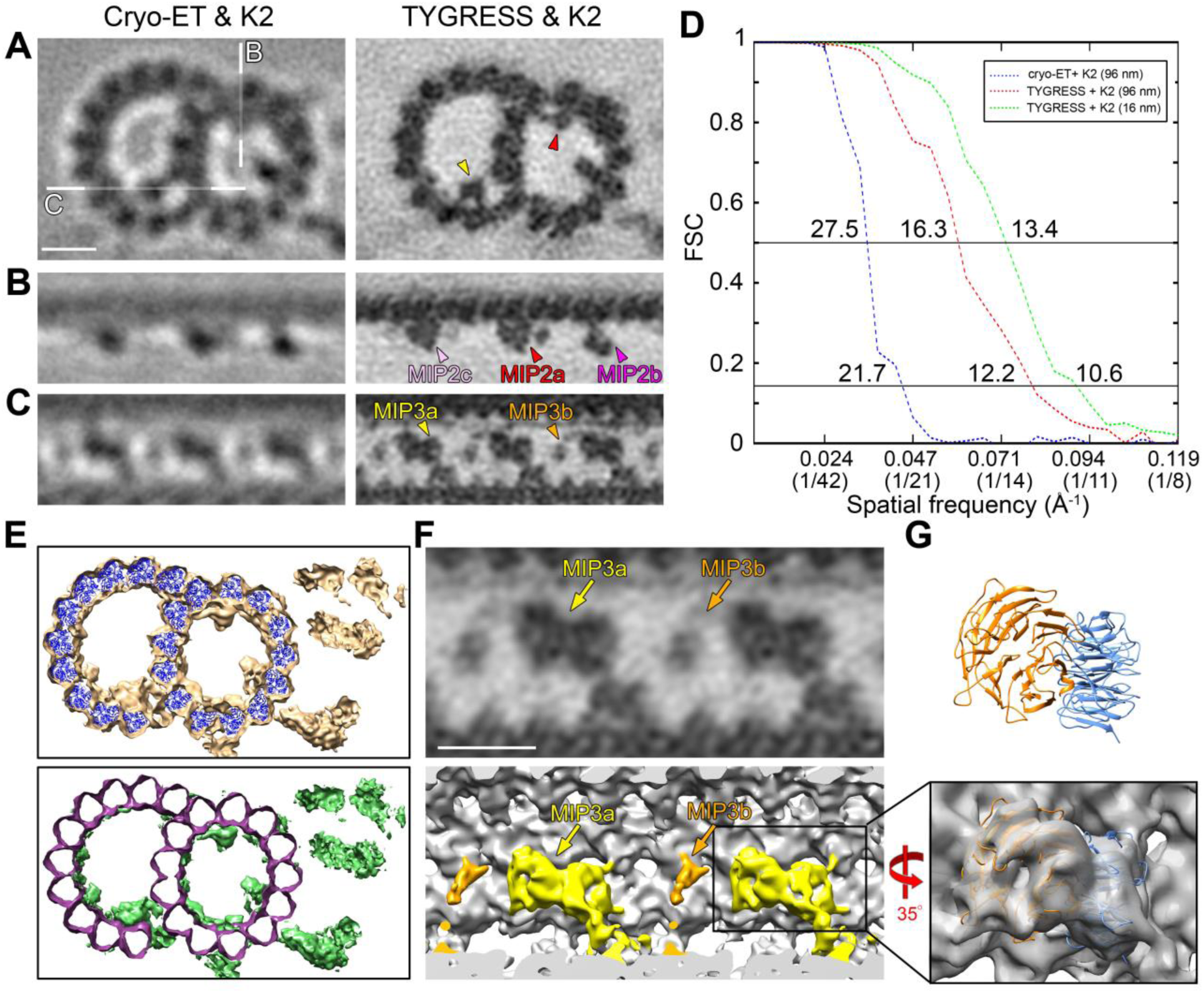
Comparison of doublet microtubules (DMTs) and microtubule inner proteins (MIPs) visualized using different methods. (**A-C**) Comparisons of EM slices of the *Tetrahymena* axonemal DMT obtained using cryo-ET and a K2 camera (left column) and TYGRESS and a K2 camera (right column). Cross-sectional (**A**; viewed from the proximal side of the axoneme) and longitudinal (**B** and **C**) EM slices; white lines in (**A**, left) indicate the locations of the longitudinal slices (**B** and **C**). Scale bar, 10 nm. (**D**) Resolution estimates using the Fourier Shell Correlation (FSC) (Harauz and Van Heel 1986) of the 96-nm axonemal repeat obtained with cryo-ET (blue) and TYGRESS (red) and additional averages of the 16-nm microtubule repeat (green) are shown; sub-averaging further improves the final resolution from 12.2 Å to 10.6 Å at the 0.143 FSC criterion (Rosenthal and Henderson 2003). (**E**) A difference map (bottom, green) was calculated between the *Tetrahymena* DMT structure obtained with TYGRESS (top, beige) and a docked pseudo-atomic model with tubulin dimers (top, blue). The remaining density (bottom, green) represents MAPs and MIPs attached to the microtubule lattice. (**F**) EM slice (top) and 3D isosurface rendering (bottom) of MIP3a (yellow) and MIP3b (orange) after averaging the 16-nm repeats. Scale bar, 10 nm. (**G**) A predicted pseudo-atomic model of FAP52 (top) was docked to our MIP3a density (bottom).

**Figure 3.**
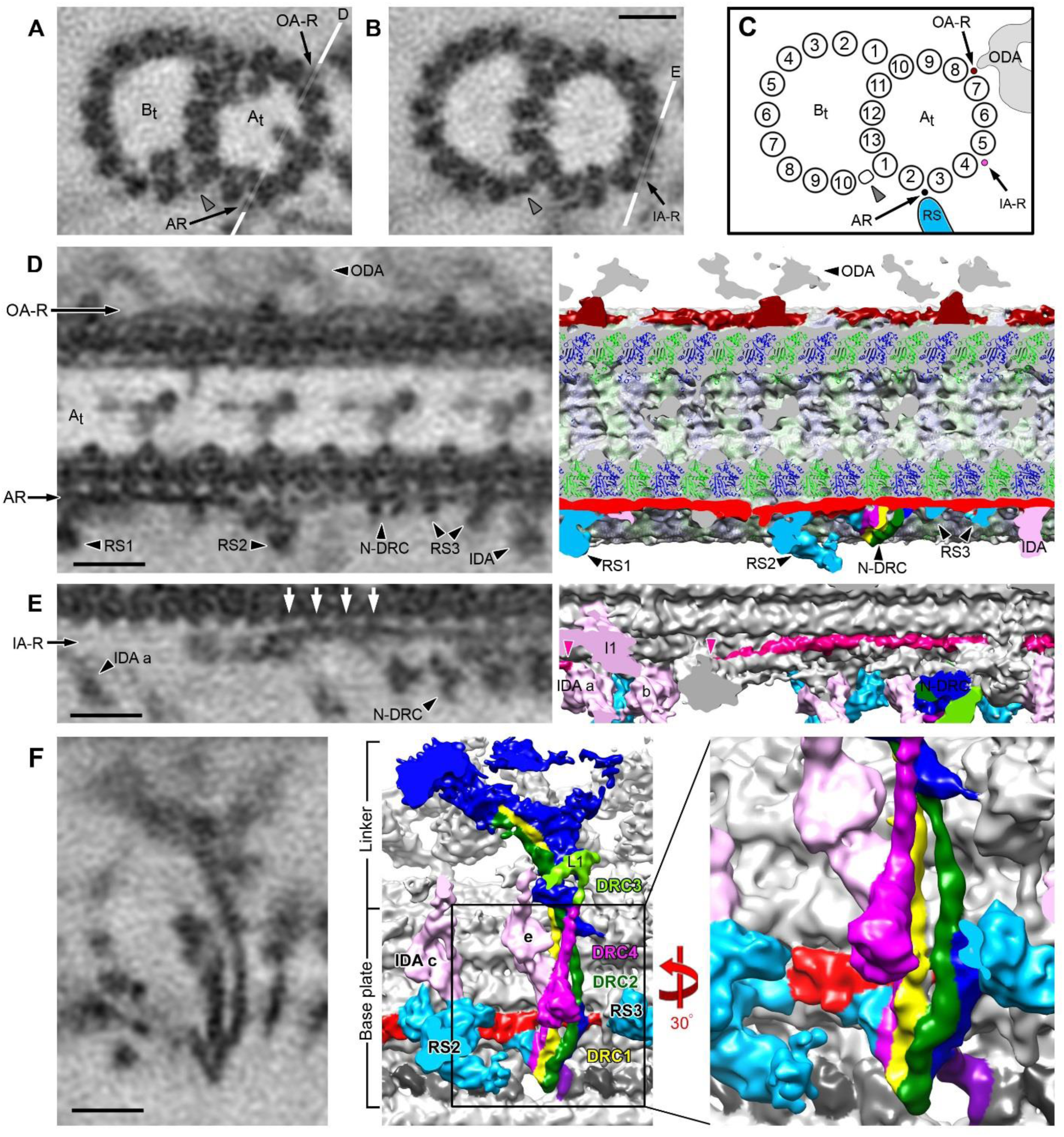
Filamentous structures and subunit architecture of the nexin-dynein regulatory complex (N-DRC) located outside the DMT were visualized by TYGRESS. (**A** and **B**) Cross-sectional EM slices of averaged 96-nm axonemal repeats in two different positions showing three well-resolved filamentous structures: the 24-nm outer dynein arm ruler (OA-R), 96-nm axonemal rulerAR, and inner dynein armIA-R. White lines indicate the locations of the EM slices shown in **D** and **E**. The inner junction (IJ) is indicated by a gray arrowhead. (**C**) Schematic diagram showing the locations of the OA-R, AR, and IA-R. The IJ is indicated by a gray arrowhead. Other labels: ODA, outer dynein arm; At/Bt, A/B-tubule; RS, radial spoke. (**D**) Longitudinal EM slice (left) and 3D isosurface rendering (right) of the 96-nm axonemal repeat showing the OA-R (dark red) and AR (red). A tubulin dimer model is fitted into the 96-nm axonemal repeat density based on the published information regarding the positions of α/β-tubulin on the DMT (Maheshwari et al. 2015). The termini of AR were well resolved. The three radial spokes (RSs 1-3, light blue), N-DRC components (purple, yellow, and dark green), and inner dynein arms (IDAs) are assembled onto the DMT through the AR. (**E**) Longitudinal EM slice (left) and 3D isosurface rendering (right) of the 96-nm axonemal repeat showing the IA-R (magenta). The IA-R starts between the RS1 and RS2 (indicated by a magenta arrowhead) and attaches to and stretches along the microtubule, connecting to the tail of inner dynein arm a (indicated by a magenta arrowhead). Scale bars, 10 nm. (**F**) EM slice (left) and 3D isosurface rendering (middle and right) of the N-DRC structure. DRCs 1-4 were well resolved. DRC 1, 2 and 4 exhibit filamentous structures, which comprise the base plate of the N-DRC and expand to the linker domain of N-DRC. DRC1, yellow; DRC2, dark green; DRC3, light green; DRC4, purple; unknown N-DRC components are shown in dark blue; the tails of inner dynein arm IDA c and e, pink; the bases of RS2 and RS3, light blue. Scale bar, 10 nm. (**F**, right) Close-up of the 3D isosurface rendering of the N-DRC structure rotated 30° from the structure in (**F**, middle). The coiled-coil strands of DRC 1, 2 and 4 are well resolved and comprise the base plate of the N-DRC, which contacts the 96-nm ruler (red) and the DMT.

**Figure 4.**
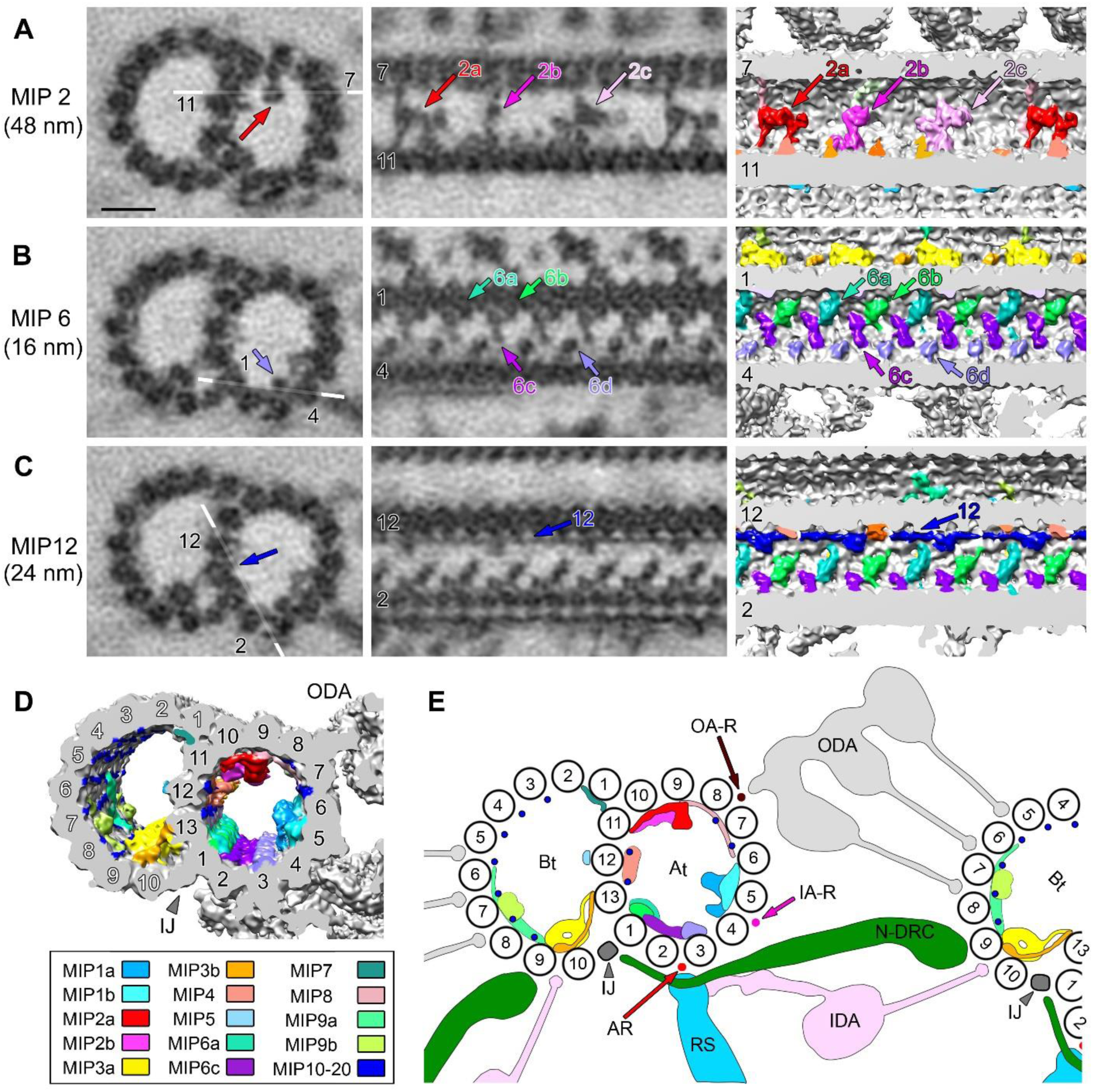
Unique structural characteristics of MIP complexes from isolated but intact *Tetrahymena* axonemes resolved by TYGRESS. (**A-C**) Cross-sectional (left column) and longitudinal (middle column) EM slices and longitudinal 3D isosurface renderings (right column) of the averaged DMT show the structural details and diversity of selected MIP complexes. White lines in the cross-sections indicate the locations of the longitudinal EM slices. The microtubule protofilament numbers of the A- and B-tubules are indicated by white and black labels, respectively. MIP periodicities are indicated by numbers inside brackets. Scale bar, 10 nm. (**D**) The 3D isosurface rendering of the 96-nm axonemal repeat shows the locations of 20 MIP complexes in the cross-sectional view. MIPs are colored and numbered according to their locations in the cross-section. The IJ is indicated by the gray arrowhead. (**E**) Schematic summary showing the locations and interactions of axonemal complexes. Positions of axonemal complexes including the microtubule protofilaments, At/Bt, IJ, AR, OA-R, IA-R, RS, IDA/ODA, N-DRC and MIPs are shown.

## Resolution improvement of the TYGRESS reconstruction

The use of direct electron detectors (e.g., K2 camera) is a key reason for the recent success and resolution improvement of SP-cryo-EM (Chiu et al. 2015). However, using cryo-ET in combination with a K2 camera for subtomogram averaging of the 96-nm repeat in intact ciliary axonemes resulted only in a small resolution improvement (27.5 Å, FSC = 0.5) over the cryo-ET average using a CCD camera [30 Å, (Lin and Nicastro 2018)] (Fig. 2, A to D). In contrast, using TYGRESS in combination with a K2 camera, we reconstructed the 96-nm repeat at up to 12 Å resolution (FSC=0.143, measured at the DMT) (Fig. 2, A to D). This is the highest resolution of the intact axoneme structure reported so far. Only SP-cryo-EM reconstructions of isolated and dialyzed DMTs have achieved a slightly better resolution (Ichikawa et al. 2017), however, with the caveat that all external and a few internal DMT structures were missing due to the isolation procedure. With the resolution of the TYGRESS reconstruction many unprecedented structural details were revealed. Individual tubulin monomers of the DMT walls can be distinguished (Fig. 2B, right panel), allowing to generate a pseudo-atomic model by fitting the tubulin dimer model derived from the high-resolution cryo-EM map of dynamic GDP state (Zhang et al. 2015) into the DMT structure. By subtracting the DMT pseudo-atomic model from the DMT density we calculated a structural difference map that reveals densities of a plethora of accessory proteins and complexes that are assembled on the DMT scaffold (Fig. 2E).

The DMT is a highly repetitive structure and by increasing the number of averaged particles from 18,731 to 112,386 by averaging the 16-nm repeating units of the DMT wall and several MIP structures, we were able to further improved the resolution from 12 Å to 10.6 Å (Fig. 2D). At this resolution, the pseudo-atomic models of individual ciliary proteins and/or domains could be reliably fitted into the structure (Fig. 2, F and G).

## Molecular rulers facilitate docking of Microtubule associated proteins (MAPs) to the outer surface of DMTs

Our resolution-improved structure of the intact axoneme enables the detailed visualization of previously seen and unknown MAP structures. Several of these MAPs are filamentous and appear to be important adaptors for binding of other accessory structures to the outer surface of DMTs (Fig. 3, A to E, and Fig. S4A). A previous study identified the FAP59/172 complex as “96-nm axonemal ruler” that is important for proper attachment of radial spokes RS1 and RS2, as well as inner dynein arms to the DMT (Oda et al. 2014). The FAP59/172 complex was roughly localized using genetics and cryo-ET of cloneable tags, but the ruler itself could not be visualized (Oda et al. 2014). In the TYGRESS average the filamentous structure of the FAP59/172 complex (“axonemal ruler” (AR)) can be clearly observed running along the DMTs in the outside cleft between protofilaments A2 and A3, with two globular domains near the bases of radial spokes RS1 and RS2 (Fig. 3, A to D, and Fig. S4A). Several axonemal complexes that have been shown to be essential for ciliary motility seem to directly connect to the FAP59/172 filament, including radial spokes RS1-3, the nexin-dynein regulatory complex (N-DRC), I1 tether/tether head (I1 T/TH), and inner dyneins b, e, g, d (Fig. 3D and Movie S2). In contrast, inner dyneins a and c are attached to the front-prong of radial spoke RS1 and RS2, respectively. This is consistent with the previous study that demonstrated that the FAP59/172 complex is critical for docking of these structures (Oda et al. 2014).

Similarly to the above mentioned “axonemal ruler” (AR), we resolve another long filamentous structure, here termed “inner dynein arm (IDA) ruler” (IA-R), running along the outer surface of the DMT in the outside cleft between protofilaments A4 and A5 (Fig. 3, B and C, and Fig. S4A). Its proximal terminus starts at a density connected with I1 between RS1 and RS2, and connects to the tail of IDA a, d, and g (Fig. 3E). The direct connections between IA-R and protofilament A4 with 4-nm periodicity are clearly resolved (Fig. 3E, white arrows). Interestingly, this periodic connection between IA-R and protofilament A4 is discontinued in the region of the I1 dynein complex (Fig. 3E). Therefore, the IA-R may play a similar role as the axonemal ruler by determining the periodic docking of several IDAs. To date, the molecular composition of IA-R has not been identified.

The outer dynein arms (ODAs) attach to the outer surface of DMTs with 24 nm periodicity, but the molecular mechanism of this regular docking remains unclear. Previously, the 24-nm-long ODA docking complex (ODA-DC) was proposed to be responsible for the assembly and 24-nm periodicity of ODAs, but the localization of the ODA-DC remained undetermined (Owa et al. 2014). However, it was recently shown that *Chlamydomonas* ODAs can assemble onto DMTs lacking ODA-DC from the *oda3* mutant *in vitro*, with proper periodicity (Oda et al. 2016). In addition, the cryo-ET structure revealed that the part of ODA-DC contacts with DMT near to the microtubule binding site of ODA, but the rest of the complex binds to the outer surface of ODA (Oda et al. 2016). Together, this suggested that ODA-DC may not be able to serve as a molecular ruler. In the TYGRESS average we observe a filamentous structure, here termed “ODA ruler (OAR)”, running along the outside cleft between protofilaments A7 and A8 (Fig. 3, A and C, and Fig. S4A). A globular density of OA-R with 24-nm periodicity seems to provide the docking site for ODAs onto the DMT (Fig. 3D).

## Subunit organization of the N-DRC base plate and linker base

The N-DRC is separated in two major regions, the base plate that is important for the N-DRC binding to the DMT, and the linker region that connects to the neighboring DMT (Fig. 3F), which is critical for both axoneme integrity and ciliary motility (Alford et al. 2016). In contrast to previous cryo-ET studies that observed the N-DRC base plate as a single rod-shaped density (Song et al. 2015), in the TYGRESS reconstruction three N-DRC subunits are well-resolved as three long filamentous structures that slowly twist around each other (shown in purple, yellow, and dark green in Fig. 3F). We proposed that these filamentous structures are DRC1, 2, and 4, subunits that have previously been localized to the N-DRC base plate (Heuser et al. 2009), and that are predicted to be enriched in coiled-coil domains (Lin et al. 2011).

Our structure reveals two similar N-DRC filaments (yellow and dark green in Fig. 3F) that reach from the inner junction between the A- and B-tubule, to the proximal lobe of the N-DRC, and directly connect to the 96-nm AR and the DMT. This suggests an important role of these subunits for the assembly and docking of the N-DRC. We propose that these filaments are DRC1 and DRC2, because it has been shown that their absence in *pf3* and *ida6* mutant axonemes causes the loss of the entire base plate (Heuser et al. 2009, Lin et al. 2011, Wirschell et al. 2013, Bower et al. 2018). In contrast, the shorter filament with associated globular domain (purple in Fig. 3F) binds on top of DRC1 and 2, and thus is likely not essential for base plate assembly, but seems to directly connect to the linker arm L1 (light green in Fig. 3F). These structural features and previously described mutant phenotypes are consistent with the short base plate filament being DRC4 that connects to DRC3 in the L1 arm (Song et al. 2015).

The N-DRC has been shown to connect adjacent DMTs (Huang et al. 1982, Nicastro et al. 2006, Heuser et al. 2009), which is critical for restricting and thus transforming interdoublet sliding into the ciliary bending motion (Summers and Gibbons 1971, Heuser et al. 2009). However, it is unclear if the Nexin-link can stretch considerably to accommodate the relative displacement between neighboring DMTs during bending (Warner 1976, Heuser et al. 2009), or if the nexin-linker can release and rebind to the neighboring DMT as it slides by (Bozkurt and Woolley 1993, Minoura et al. 1999, Heuser et al. 2009). The filamentous and rope-like nature of the DRC1/2/4 complex likely provides high tensile strength with some lateral flexibility to the N-DRC, as described for other filamentous arrangements like intermediate filaments or collagen (Sun et al. 2002, Mucke et al. 2004). The tensile strength could be important for N-DRC function while resisting considerable mechanical stress during ciliary beating. The stability against “pulling” forces that would separate neighboring doublets from each other is also consistent with the importance of the N-DRC for maintaining the axoneme integrity (Alford et al. 2016). Consistent with this function, lack of either one of the three subunits DRC1, 2 and 4 leads to severe motility defects and is linked to the ciliary disease Primary Ciliary Dyskinesia (PCD) (Austin-Tse et al. 2013, Horani et al. 2013, Wirschell et al. 2013, Olbrich et al. 2015). Lastly, the structural architecture seems inconsistent with a nexin-linker that could stretch considerably during bending, and favors an N-DRC that would release and rebind repeatedly during ciliary beating.

## Inner junction

The inner junction (IJ) between the A- and B-tubule (also called the “B-11th density”) (Linck et al. 2014), was previously shown to be a non-tubulin connection between protofilaments A1 and B10 (Nicastro et al. 2011). Our improved structure provides additional detail, suggesting that the inner junction consists of two similar-sized subunits that alternate along the DMT length with 8 nm periodicity of the heterodimer (Fig. S4, B to G, gray arrowheads). At the previously reported hole on the inner junction with 96-nm periodicity, one of these subunits is missing, likely due to interference with the C-terminal domains of DRC1 and 2 (Fig. S4, C and F, black arrowheads).

Each of the two IJ subunits has an estimated molecular weight of ~20 kDa. This lends support to a previous cryo-ET study that located BCCP-tagged FAP20 (20 kDa) in or near the inner junction in *Chlamydomonas* flagella (Yanagisawa et al. 2014). Interestingly, mutants lacking FAP20 show severe defects in the flagellar motility (Yanagisawa et al. 2014).

Although the second IJ subunit has not yet been identified, the detailed structural map provided by TYGRESS will greatly aid future localization studies of axonemal components, or even the docking of high-resolution structures and models. All above described MAPs and the inner junction were completely missing from the high-resolution SP-cryo-EM structure of isolated DMTs (Ichikawa et al. 2017), highlighting the critical need for a method like TYGRESS that can visualize cellular structures *in situ* at high resolution.

## A plethora of MIPs binds to the inner surface of the of A- and B-tubule walls

MIPs were discovered as structures that bind periodically to the inner surface of the ciliary DMT walls (Nicastro et al. 2006). However, since then MIPs have also been described for other hyper-stable microtubule populations, such as the ventral disc microtubules in Giardia that are important for the pathogen’s suction-cup-like attachment to the intestine (Schwartz et al. 2012), or the subpellicular microtubules that provide mechanical stability to apicomplexan parasites (e.g. Plasmodium and Toxoplasma) (Cyrklaff et al. 2007). Although MIPs are proposed to be important for increasing the stability of DMTs, functional studies have not been possible, because MIP proteins have not yet been identified. A recent SP-cryo-EM study of isolated DMTs has visualized the ciliary MIPs with considerably increased resolution (Ichikawa et al. 2017). However, the study has two limitations: a) the DMTs adopt a preferred orientation in the thin ice layer on the EM grids, which has previously been observed (Sui and Downing 2006) and resulted in anisotropic resolution (similar to a missing wedge in cryo-ET); and b) the isolation procedure of the DMTs involved high salt extraction and dialysis to remove MAPs from the outside surface of the DMT; however, this sample preparation also resulted in the dissociation of other structures, such as the inner junction proteins (Ichikawa et al. 2017).

In the TYGRESS reconstruction, the MIPs were well resolved and can be grouped into 20 discrete densities (MIPs 1-20) (Fig. 4, and Figs. S5 and S6). MIPs 1-6, which have been previously observed (Nicastro et al. 2006, Maheshwari et al. 2015), were revealed with greater structural detail, allowing to distinguish MIP substructures. For example, the three substructures of MIP2 (MIP2a, b, and c), previously appeared as three similar globular densities by cryo-ET (Fig. 2B, left panel). In contrast, the TYGRESS average clearly resolves structural differences between MIP2a, b, and c (Fig. 2B, right panel, and Fig. S5). Thus, the periodicity of the MIP2 proteins has to be corrected from the previously reported 16 nm to 48 nm. Similarly, MIP6, which was reported as a continuous structure spanning protofilaments A1-3, is now resolved as four discrete substructures MIP6a-d (Fig. 4B, and Fig. S5).

MIPs 7-20 were not visualized by previous cryo-ET and subtomogram averaging studies but are now resolved by TYGRESS of intact ciliary axonemes (Fig. 4C, Figs. S5 and S6), and by SP-cryo-EM of isolated DMTs (Ichikawa et al. 2017). MIP7 binds to protofilaments A11and B2 along the MT axis with 16 nm periodicity (Fig. S5). MIP8 and MIP9 are resolved for the first time, which run perpendicular to the microtubule axis and span protofilaments A6-9 and protofilaments B8-10, respectively, with 48 nm periodicity (Fig. S5). Remarkably, MIPs 10-20 are filamentous MIPs that follow individual protofilaments along the DMT axis (Fig. 4C and Fig. S6). These MIPs are considerably thinner than the 96-nm axonemal ruler. Previous studies found that the hyperstable ribbon of three adjoining protofilaments contains tektin filament (with tektins A-C) among other accessory proteins (Linck et al. 2014). We showed previously that the hyperstable ribbon forms the DMT midpartition, i.e. protofilaments A1 and A11-13 (Linck et al. 2014). Therefore, MIP12 (Fig. 4C and Fig. S6) and/or MIP13, which are attached to protofilaments A13 and A12, respectively, may be the tektin filaments. Interestingly, we also find that some MIPs are periodically interconnected with adjacent MIPs, such as the connected filamentous MIPs 17 and 18, as well as MIPs 19 and 20, or connections between non-filamentous and filamentous MIPs, such as MIPs 4a-e connecting with MIPs 12 and 13 every 48 nm, MIPs 8a-b connecting with MIPs 10-11 every 48 nm, and MIPs 9a-b connecting with MIPs 17-20 every 48 nm (Figs. S5 and S6). This architecture suggests that the MIPs form a complex network that could considerably increase the stability of the DMTs, especially by strengthening the usually relatively weak lateral protofilament-to-protofilament interactions of microtubules. The axoneme is an important scaffold in cilia that serves as a persistent platform for the attachment of hundreds of accessory proteins, making the stabilizing MIP network essential for maintaining DMT integrity under the considerable mechanical stress during ciliary beating.

In summary, the results not only provide a better understanding of the functional organization of the DMTs and ciliary axoneme (Fig. 4, D and E), but they also demonstrate that TYGRESS can resolve macromolecular complexes at unprecedented resolution while they are maintained *in situ*, i.e. in their cellular context. Here, we have used it to reconstruct the 96-nm axonemal repeat at 12 Å. We expect the resolution to improve further with energy-filtered data recorded on more stable cryo stages, and using patch-based motion correction (MotionCor2) (Zheng et al. 2017) and CTF correction to take into account the variable high of particles in a sample (Turonova et al. 2017). Furthermore, given larger datasets, single-particle classification could be used to calculate more homogeneous class averages with density features. Generally, the TYGRESS method should improve the resolution of cellular 3D reconstructions that are amenable to subtomogram averaging, especially in cases where sample thickness and radiation damage are the main resolution-limiting factors. TYGRESS is ultimately a SP-reconstruction method and thus will continue to benefit from the same future advances that improve SP-cryo-EM of isolated particles. However, similar to SP-cryo-EM, preferred orientation of structures, e.g. focal adhesion complexes (Geiger et al. 2009) that are always oriented parallel to the EM grid, cannot be overcome by TYGRESS unless tilted HD images are included in the reconstruction. An added advantage of the cryo-ET and subtomogram averages is that they can be used to classify structurally heterogeneous complexes in 3D before calculating TYGRESS class averages. One current challenge with TYGRESS is that data acquisition and image processing is time-consuming, but future developments of high-throughput tilt-series acquisition, possibly in minutes using continuous camera exposure while the sample is tilted in quick increments with highly eucentric TEM specimen stages, and continued improvements to automate tilt series alignments and tomogram reconstructions, could considerably decrease the time needed for TYGRESS data acquisition and processing.

## Acknowledgements

We thank Chen Xu for training and maintaining the electron microscopy facility at Brandeis University. We thank Alexis Rohou (Janelia Research Campus), John Heumann and David Mastronarde (Univ. of Colorado in Boulder) for technical advising and helpful discussion about image processing. We thank Thomas Ni (UT Southwestern) for critically reading the manuscript. We thank Rui Zhang at Washington University for providing the EM structure of tubulin dimer for docking.

## Author contributions

D.N. designed experiments. Z.S. and X.F. programmed, K.S. collected data and processed the data with Z.S.. N.G. contributed significant scientific and technical insights throughout the project. Z.S., K.S., X.L., N.G., and D.N. wrote the manuscript. All authors contributed to discussions and revisions of the manuscript.

## Materials, instrumentation, and data acquisition

### Cryo-sample preparation

Wild type *Tetrahymena thermophile* axonemes were isolated from strain CU428 cells, as previously described (Lin et al. 2014). Briefly, cilia were detached from the cells using the pH-shock method (Witman et al. 1972) and purified by centrifugation at 2,400 *g*, 4 °C, for 10 min (twice). Purified cilia were demembranated using 1% IGEPAL CA-630 (Sigma-Aldrich) in HMEEK buffer (30 mM HEPES, pH 7.4, 5 mM MgSO_4_, 1 mM EGTA, 0.1 mM EDTA, and 25 mM KCl) and axonemes were collected by centrifugation at 10,000 *g*, 4 °C, for 10 min. The axoneme pellet was carefully resuspended in HMEEK buffer and stored on ice for cryo-sample preparation within 4 hours.

Cryo-samples were prepared using previously described methods (Lin et al. 2014). Briefly, Quantifoil grids (Quantifoil MicroTools GmbH, Germany) were glow discharged for 30 s at −40 mA before use, coated with 10 nm colloidal gold (Sigma–Aldrich), and loaded on a plunge freezing device. Then, 3 μl of axoneme sample and 1 μl of a five-fold-concentrated 10-nm colloidal gold solution (Iancu et al. 2006) were added to the grid and mixed within seconds. The grid was blotted with filter paper for 1.5 –2.5 s and immediately frozen by plunging into liquid ethane. The vitrified samples were then stored in liquid nitrogen until examined by electron microscopy.

### Image acquisition

The frozen grid was mounted in a cryo-holder (Gatan Inc., Pleasanton, CA) and imaged on a Tecnai F30 transmission electron microscope (FEI, Inc., Hillsboro, OR) equipped with a field emission gun and operated at 300 keV. The data were collected under low-dose conditions using the SerialEM software (Mastronarde 2005). For each intact axoneme, two sets of data were collected using the K2 direct detector (Gatan Inc., Pleasanton, CA) at a magnification of 9400×. First, a movie stack (80 frames) was collected at 0° with a total electron dose of ~30 e/Å^2^ (HD image) at varying defocuses of −1.5 to −3 μm in the K2 super-resolution mode using; then, a typical tilt series with an accumulated electron dose of ~100 e/Å^2^ (LD images) was recorded at a defocus of −8 μm using a tilting range from −64° to 64° in 2.0° increments in the K2 counting mode. At each tilt angle, a movie stack (5 frames) with an exposure time of 2 second and election dose of 1.5 e/Å^2^ was recorded. The resulting pixel sizes of the 0° HD image and the LD images were 0.2112 nm and 0.4224 nm, respectively. The parameters used for data collection are summarized in Table S2.

### Image processing

Full-frame motion correction of the movie stacks was performed using IMOD scripts (Kremer et al. 1996). Then, both the HD image and LD images were aligned using fiducial markers. The LD images alone were further reconstructed into a 3D tomogram by weighted back-projection using the IMOD software package. Subtomograms containing the 96-nm axonemal repeats were extracted from the tomograms, aligned, and averaged using PEET (Nicastro et al. 2006) with cross-correlation searching. The HD images were used to reconstruct high resolution structures with the TYGRESS method developed in this study (SOM text). FREALIGN (Grigorieff 2007) was used for the final reconstructions as part of TYGRESS. CTFFIND3 (Mindell and Grigorieff 2003) was used to detect defocus values. BFACTOR (Fernandez et al. 2008) was used to sharpen and filter the final reconstruction. UCSF Chimera (Pettersen et al. 2004) was used for 3D visualization and x-ray structure fitting. The pseudo-atomic models of MIP3a/FAP52 were generated and calculated from the protein sequences using the ExPASy online tool, SWISS-MODEL (Biasini et al. 2014).

### Resolution measurement

The resolution of the TYGRESS reconstruction was measured using the FSC (Harauz and Van Heel 1986). Briefly, the dataset was divided into two halves using even and odd indexes, then the two halves were reconstructed independently and the FSC between the two reconstructions was calculated.

### SOM text: TYGRESS image processing

TYGRESS is essentially a single-particle reconstruction procedure using coordinate information provided by cryo-ET to enable particle picking (Fig. 1). During imaging, two datasets were acquired for each region of interest: a 2D image (HD image) at 0° for the final TYGRESS reconstruction using an electron dose typical for conventional SP-cryo-EM, and a traditional low-dose tilt series (LD images) that was used for positional information. To minimize the effects of radiation damage in the final reconstruction, the HD image was recorded prior to the LD images (Fig. 1A).

### Alignment of the combined tilt series and tomogram reconstruction with subtomogram averaging

During image processing, each HD image was inserted into the corresponding tilt series at the angle corresponding to where the HD image was taken (for example, an HD image recorded at 0° was inserted just before the LD image at 0°) using the command ‘newstack’ in IMOD, resulting in a “combined tilt series”. The combination of HD and LD images ensured a common reference frame for the later steps in the TYGRESS procedure. The combined tilt series was then aligned based on the 10-nm gold fiducial markers using the IMOD software package. However, only the LD images were used to calculate a tomogram after alignment, and then subtomogram averaging was performed in PEET (Fig. 1B). In this work, the 96-nm repeat unit of the axoneme can be readily identified based upon structural features such as the doublet microtubule (MT) walls and the radial spokes, which allowed us to pick particles of 240^∗^240^∗^240 pixels (240^∗^0.4224 nm = 101 nm) from the noisy tomogram. The axoneme’s 9-folds symmetry facilitated determination of the initial orientation of each picked particle. Subtomogram averaging was performed using the raw tomogram and a reference, which was constantly updated using the average structure of the last iteration. In total, 18,857 particles were picked from 152 axoneme tomograms and aligned in PEET.

The HD image was excluded for tomogram reconstruction and subtomogram averaging to avoid (1) reconstruction artifacts due to uneven weighting of the HD and LD images in the tomogram, and (2) alignment bias in the initial reconstruction calculated from the HD image.

### Retrieval of coordinates and orientations for 2D SP particle on HD image

Because the HD image was aligned together with the LD images during tomographic reconstruction, we could retrieve the 2D HD image particle coordinates and orientations corresponding to the particles in the 3D tomogram. The final coordinates (*x*,*y*) and orientations (*α*, *β*, *γ*) for each 2D HD image particle are given as:

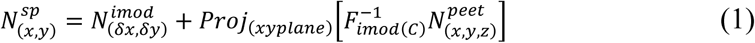

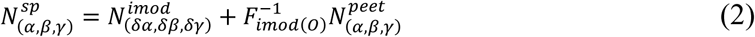

Where, (*δx*, *δy*) and (*δa*, *δβ*, *γy*) are the HD image shift and rotation from the fiducial gold alignment of the combined tilt series; 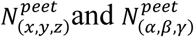 are the 3D particle coordinates and orientations in the tomogram after correction using PEET; 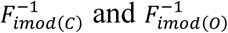 represent the inverse transformations for coordinates and orientations of particles in the HD image, which shares the same parameters as the 0° tilt LD image of the tomogram. *Proj*_(*xyplane*)_ converts the 3D coordinates to 2D coordinates in the HD image.

### 2D Particle picking from HD image

The command ‘EXCISE’ from the IMOD package was used to extract particles from the HD image according to the coordinates determined from the tomogram. In this work, the particle box size was set to 320^∗^320 pixels, which corresponds to the size of the 3D volume extracted from the tomogram. This size (320^∗^0.4224 = 135 nm) is sufficient to cover an entire axonemal repeat (96 nm). An example of how TYGRESS successfully identified the particle center from the overlapping projection images of an axoneme is shown in Fig. S1.

### Constrained Alignment of 2D HD image particles

The particle coordinates and orientations in the HD image were initialized using the results from subtomogram averaging. The subtomogram average obtained from the tomogram was used as the reference for one round of particle alignment. The low resolution of this initial reference, as well as overlap of other parts of the sample (e.g. other particles) impeded a simple alignment of particles in the HD image to this reference. To proceed, we developed an algorithm to perform the alignment of particles in images containing superimposed sample features. That is, 1) a reference of the target particle (the particle to be aligned and averaged) and the background (density corresponding to other nearby particles) are calculated according to the particle positions in Fig. 1B; 2) the two references are combined into a single reference; 3) the target particle in the HD image is aligned against the reference, constrained by shift and angular search range to not deviate from their initial subtomogram alignments by more than 4 nm in their x, y coordinates and 5 degree in their orientation (Fig. 1D) to reduce errors; 4) during constrained alignment, the background particles are fixed and only the parameters of target particle is adjusted. The constrained alignment is then repeated for all other particles in turn. A second round of alignment uses the updated particle parameters to calculate the background, and tighter constraints of 1 nm and 2 degree deviations from the parameters obtained in the previous round. After several iterations, a final set of parameters is used for 3D reconstruction. The reason for including the background particles in the combined reference is to account for all overlapping density, thereby improving the alignment accuracy of the target particle. Although this was not done in this work, the background features could also include “non-particles” that will never be aligned but represent other density in the tomogram that does not represent particles.

CTF estimation of the HD images, as well as the first round of constrained alignment, were performed using movie averages that contained all frames (1-80). To take into account local beam-induced sample motion that was not corrected by the full-frame alignment in IMOD, frame subaverages were calculated using frames 3-21, 22-40, 41-59, and 60-78 (the initial two frames were excluded due to in-frame blurring), and the second round of alignment was done using these subaverages. The aligned subaverages were then recombined in the 3D reconstruction (Fig. S2). To improve the alignment of each particle, the more variable density outside the microtubule doublet in the reference was masked.

## Supplemental figure legends

**Figure S1.**
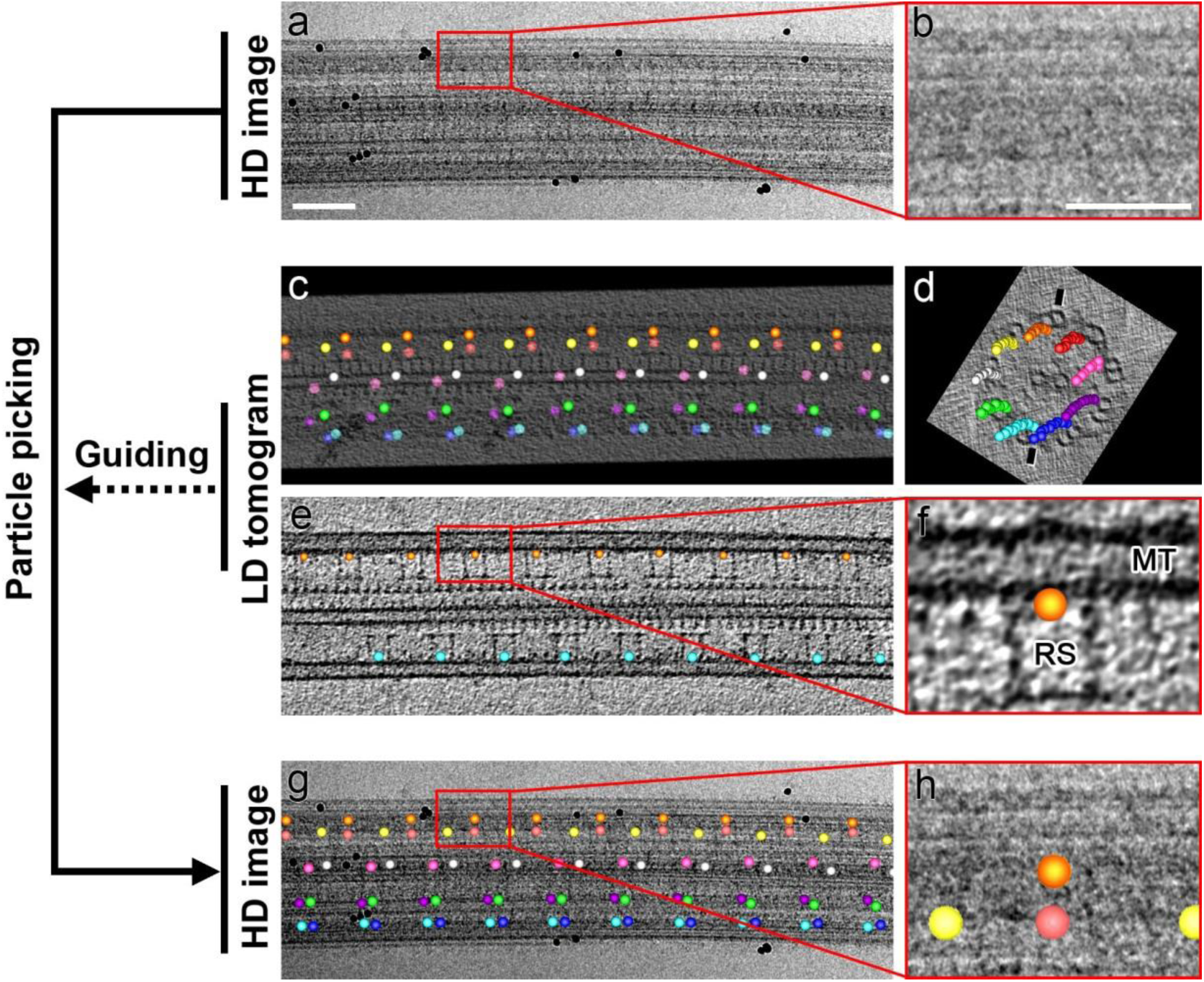
Particle picking from HD image, guided by LD-tomogram. A typical HD image (**A**) and one of its particles (**B**) show no clear features to enable particle picking because of the structural overlapping. (**C**) In the corresponding tomogram slice, many prominent particle features, such as radial spokes and microtubule walls (‘RS’ and ‘MT’ in **F**), are well-defined to help pick the same particle (yellow-red dot in **F**) in 3D (red box area in **E**). (**C-D**) show the locations for all picked particles as colored dots. Each color represents one of the 9 doublet microtubules. After the conversion of 3D coordinates into 2D, all particles can be picked out from the HD image (**G**). (**H**) The particle picked is centered at the upper yellow-red dot. Axonemes of wild type *Chlamydomonas reinhardtii* were isolated according to previously published paper (Song et al. 2015) (**A, C, E**, and **G)** scale bar, 100 nm; (**B, F**, and **H**) scale bar, 50 nm.

**Figure S2.**
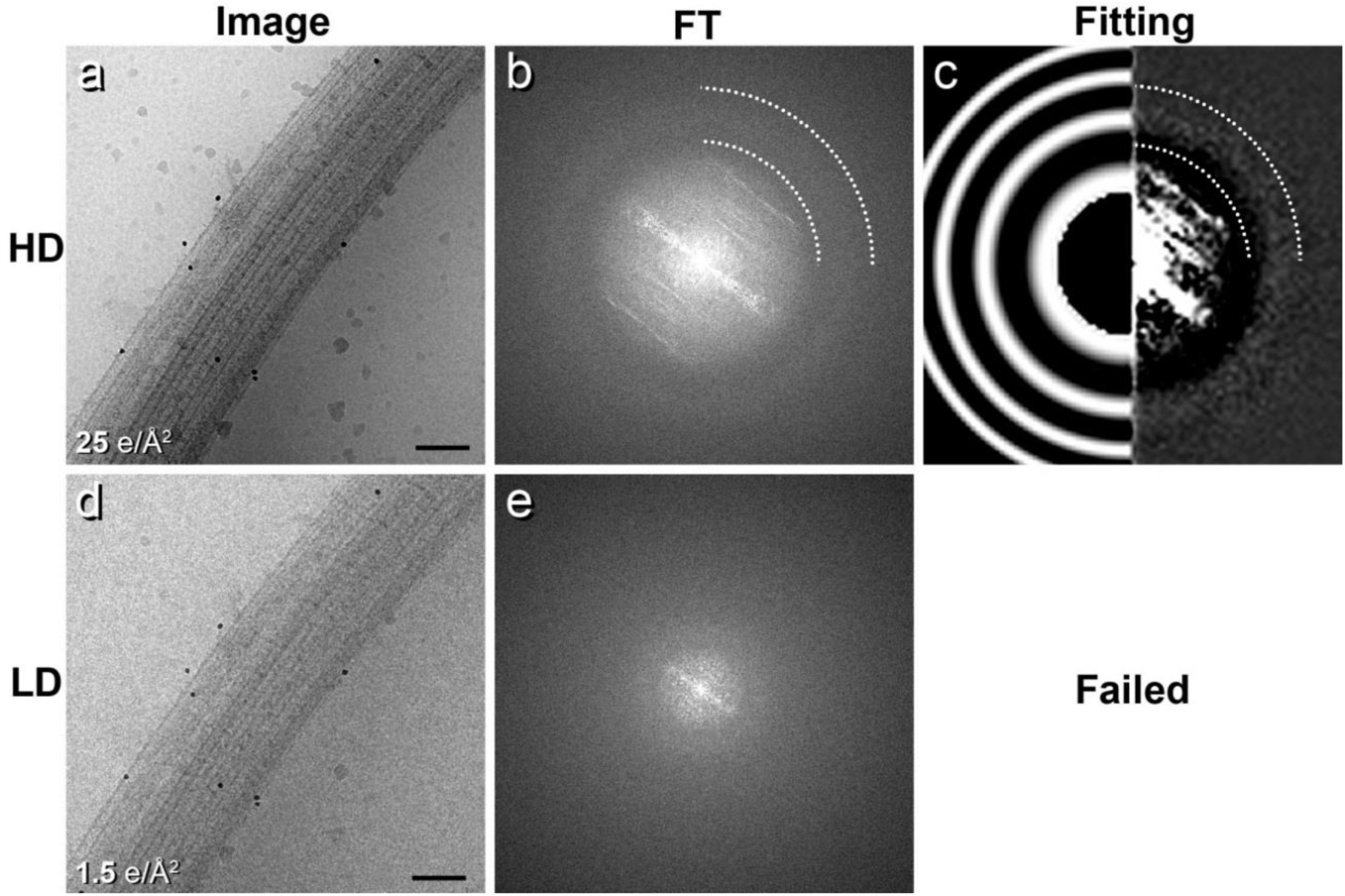
Detection of defocus values from TYGRESS HD image and LD image. A HD image from a *Chlamydomonas reinhardtii* axoneme sample (**A**), its Fourier transform (FT) (**B**), and its averaged power spectrum (**C**, right) fitting to the theoretical Thon rings (**C**, left). Strong layer lines diffracted from the repeating structures of the axoneme and two dark Thon rings (indicated by dashed lines outlining a quarter of two different circles) are visible in (**B**), but not visible in the LD image of the same sample (**D** and **E**). This caused defocus detection to fail for the LD image. The data were collected on Tecnai F30 equipped with a post-column energy filter (Gatan) and a 2k×2k charge-coupled device camera (Gatan) with a pixel size of 0.5562 nm. (**A** and **D**) scale bar, 100 nm.

**Figure S3.**
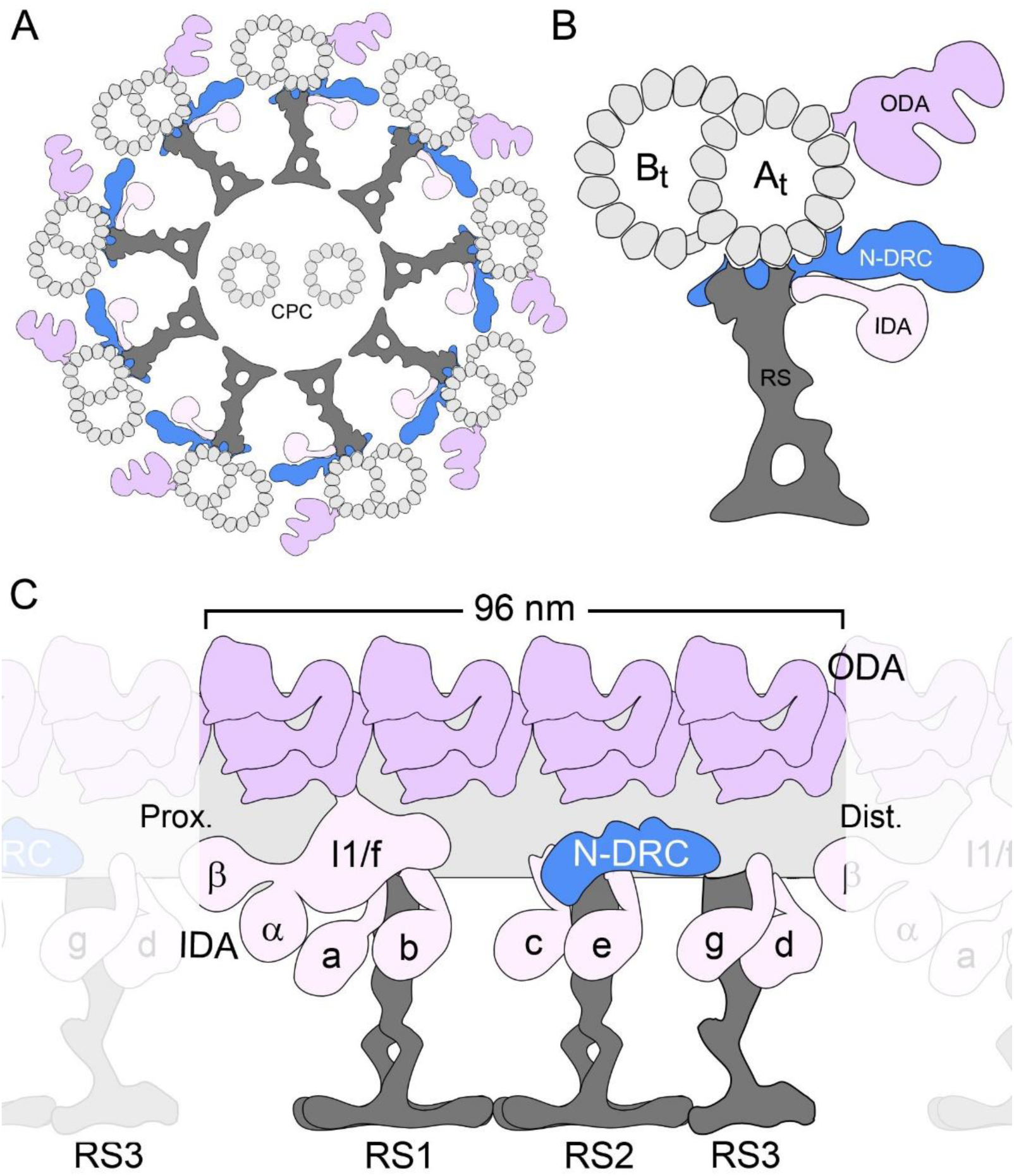
Schematic diagram of structure of axoneme. (**A**) Diagram of intact axoneme viewed in cross-section. The N-DRC is attached to the DMT by one end with it’s the other end contact to a neighbor DMT. (**B**) Diagram of an outer doublet microtubule (DMT) viewed in cross-section. (**C**) Diagram of a 96-nm-long axonemal unit that repeats along the doublet microtubule; each repeat unit contains: four outer dynein arms (ODAs), six single-headed inner dynein arms (IDAs: a, b, c, d, e and g, based on the nomenclature of *Chlamydomonas*), and one double-headed IDA (I1 or IDA f) anchored to the A-tubule (A_t_) of the doublet microtubule. Other labels: B-tubule (B_t_), central pair complex (CPC), nexin-dynein regulatory complex (N-DRC), radial spoke (RS), and microtubule polarity.

**Figure S4.**
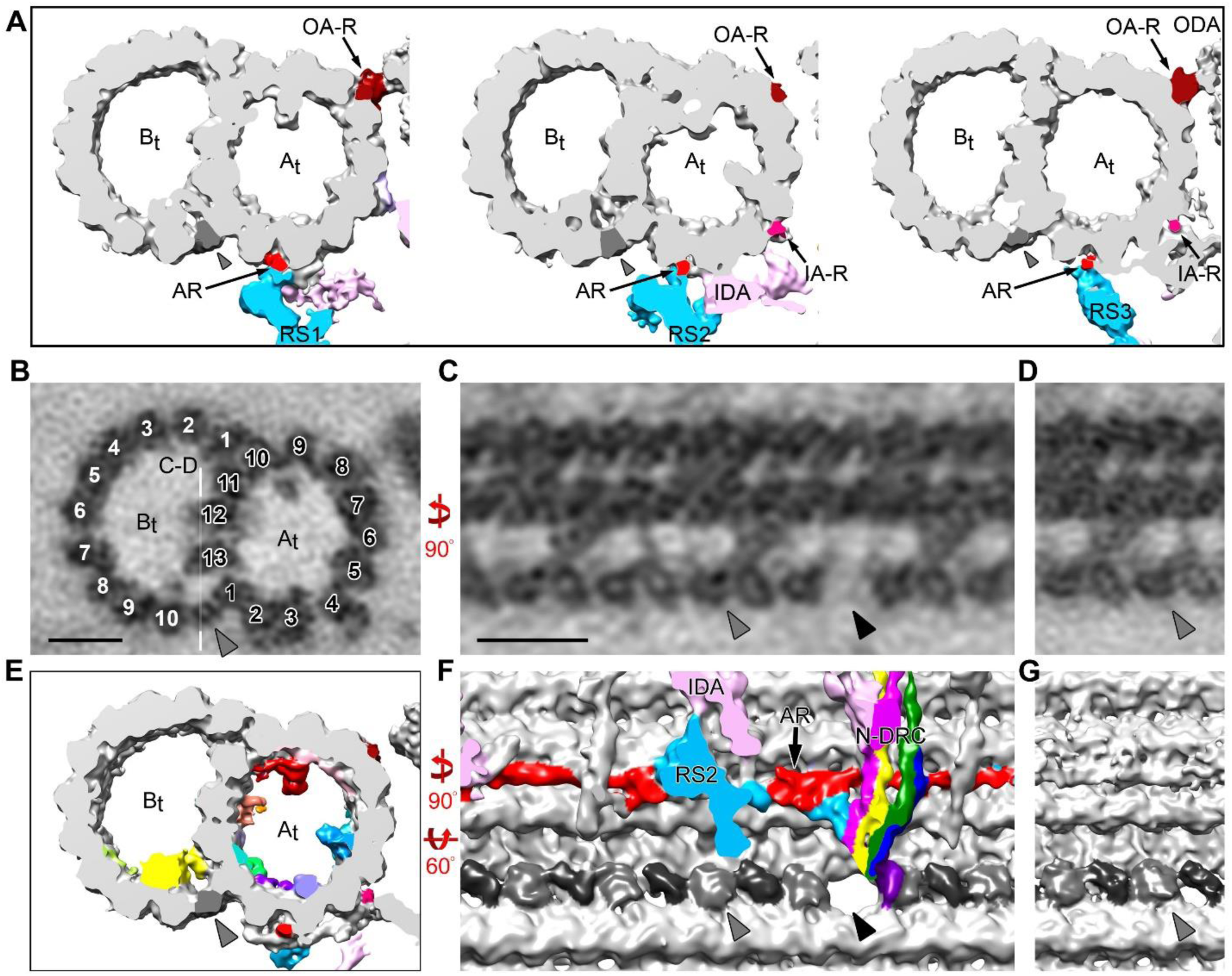
Filamentous structures outside the DMT and the inner junction (IJ). (**A**) Cross-sectional slices of isosurface renderings of the 96-nm axonemal repeat from three different locations showing the locations of the OA-R (dark red), 96-nm ruler (AR, red), and IA-R (magenta) as well as the interactions between RSs 1-3 (light blue) and the 96-nm ruler. (**B-D**) EM slices and (**E-G**) 3D isosurface renderings of the reconstructed 96-nm (**B, C, E** and **F**) and 16-nm (**D** and **G**) axonemal repeats showing the IJ structure obtained using TYGRESS. The white line in (**B**) indicates the location of the EM slices shown in (**C** and **D**). The IJ positions are indicated by the gray arrowheads. The IJ is composed of blobs (light gray and dark gray in **F** and **G**) with an approximate diameter of 4 nm. Every other blob is structurally identical. The two neighboring densities are also highly structurally similar but lie in different orientations and may therefore be the same protein. The microtubule PF numbers in the A- and B-tubules are indicated by black and white labels in (**B**), respectively. The hole in the IJ is indicated by the black arrowhead. Scale bars, 10 nm.

**Figure S5.**
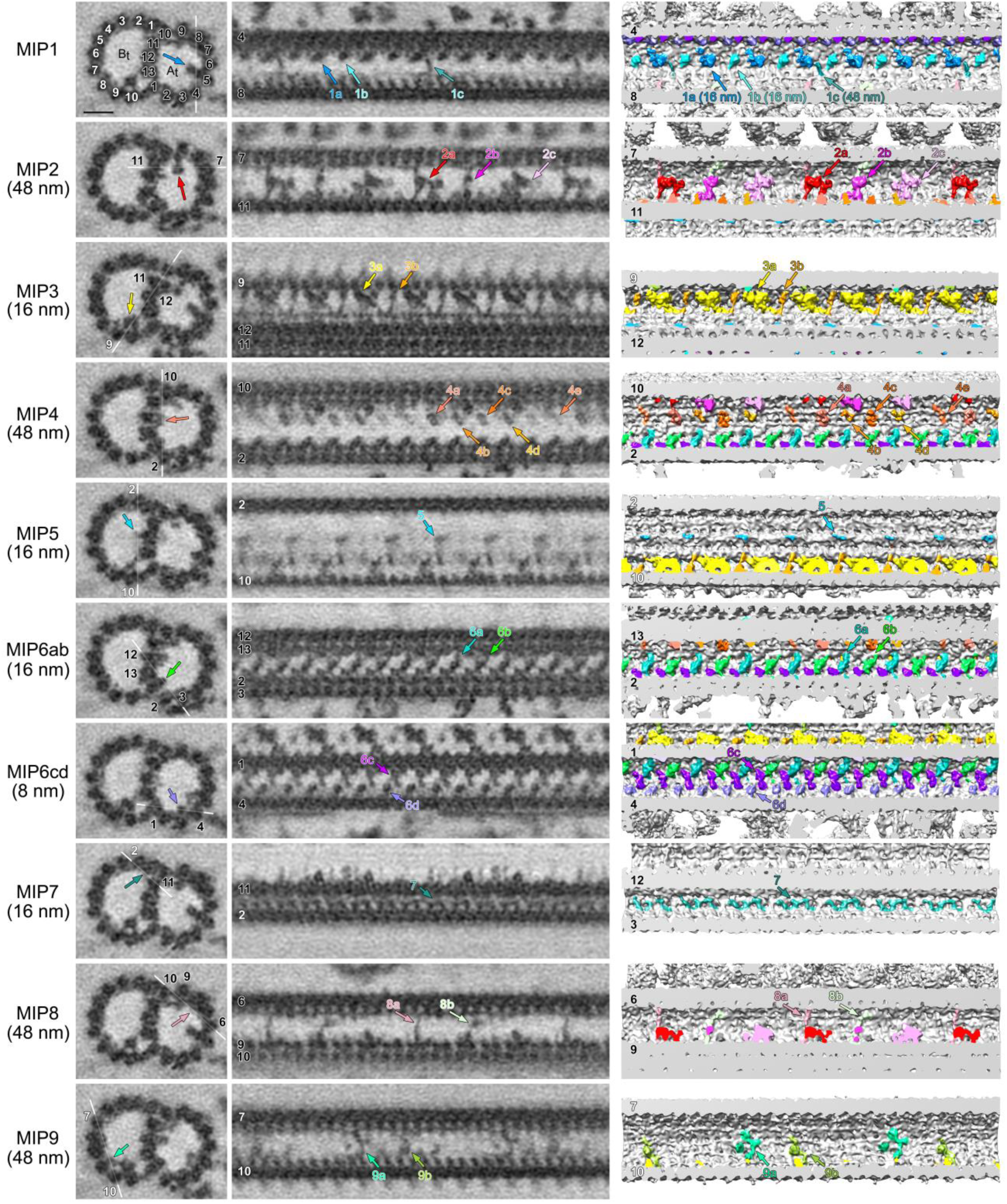
Structural characteristics of MIPs 1-9 in intact axonemes resolved using TYGRESS. Cross-sectional (left column) and longitudinal (middle column) EM slices and longitudinal views of 3D isosurface renderings (right column) of the 96-nm axonemal repeat show MIPs 1-9. The MIPs are colored and numbered according to their locations in the cross-section. MIPs present at similar locations in the cross-sectional view but in various locations in longitudinal views are further distinguished by letters. MIP periodicities are indicated by numbers inside brackets. White lines indicate the locations of the EM slices shown in the middle column. The MT protofilaments numbers in A- and B-tubules are indicated by black and white labels, respectively. Scale bars, 10 nm.

**Figure S6.**
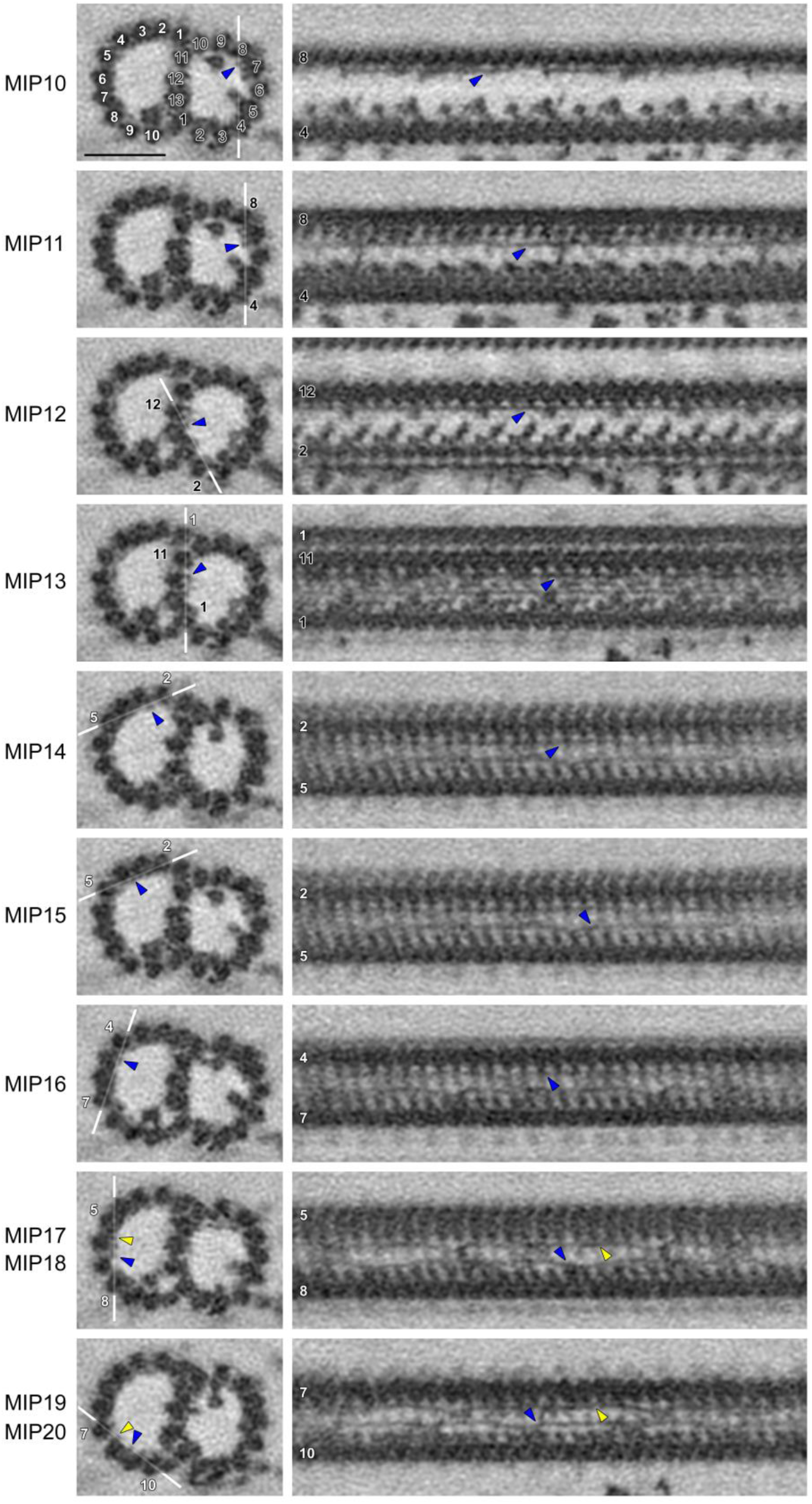
Filamentous MIPs in intact axonemes resolved using TYGRESS. Cross-sectional (left column) and longitudinal (middle column) EM slices of the 96-nm axonemal repeat show the 11 resolved filamentous MIPs. The MT protofilaments numbers in the A- and B-tubules are indicated by black and white labels, respectively. The dark blue arrows highlight the corresponding MIPs. White lines show the locations of the corresponding longitudinal EM slices. Scale bars, 20 nm.

**Table S1.**
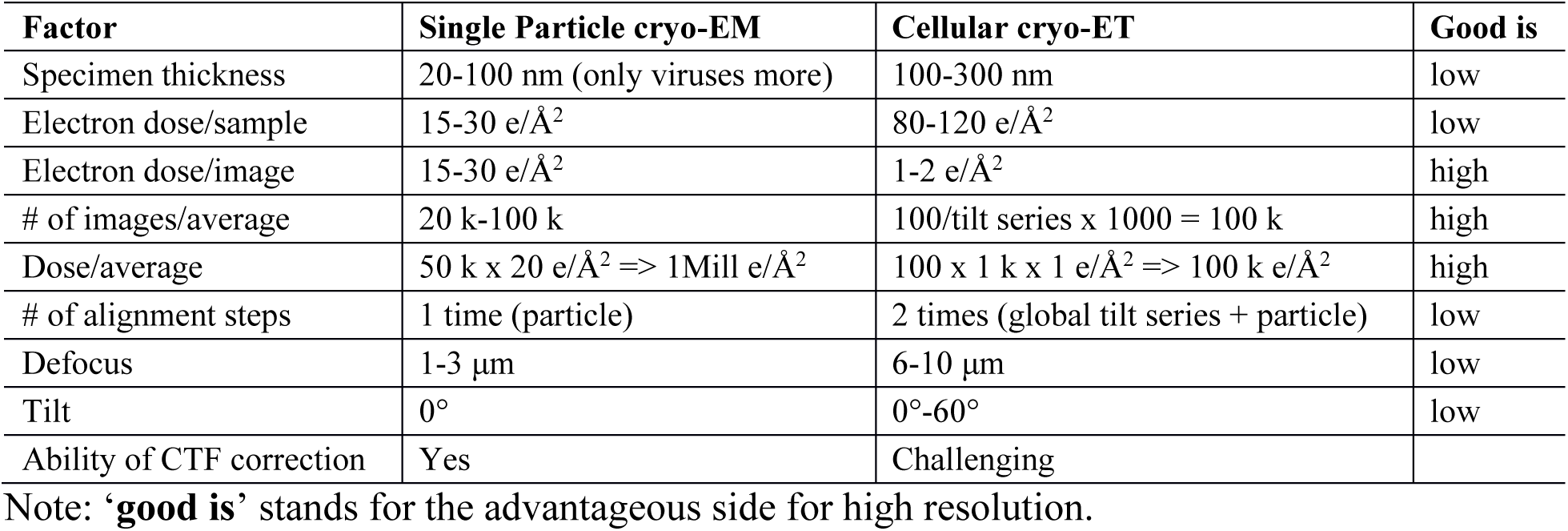
Comparison of factors that limit the resolution of cellular cryo-ET and single-particle cryo-EM.

| Factor | Single Particle cryo-EM | Cellular cryo-ET | Good is |
| --- | --- | --- | --- |
| Specimen thickness | 20-100 nm (only viruses more) | 100-300 nm | low |
| Electron dose/sample | 15-30 e/Å<sup>2</sup> | 80-120 e/Å<sup>2</sup> | low |
| Electron dose/image | 15-30 e/Å<sup>2</sup> | 1-2 e/Å<sup>2</sup> | high |
| # of images/average | 20 k-100 k | 100/tilt series x 1000 = 100 k | high |
| Dose/average | 50 k x 20 e/Å<sup>2</sup> => 1Mill e/Å<sup>2</sup> | 100 x 1 k x 1 e/Å<sup>2</sup> => 100 k e/Å<sup>2</sup> | high |
| # of alignment steps | 1 time (particle) | 2 times (global tilt series + particle) | low |
| Defocus | 1-3 μm | 6-10 μm | low |
| Tilt | 0° | 0°-60° | low |
| Ability of CTF correction | Yes | Challenging |  |

Note: '**good is**' stands for the advantageous side for high resolution.

**Table S2.**
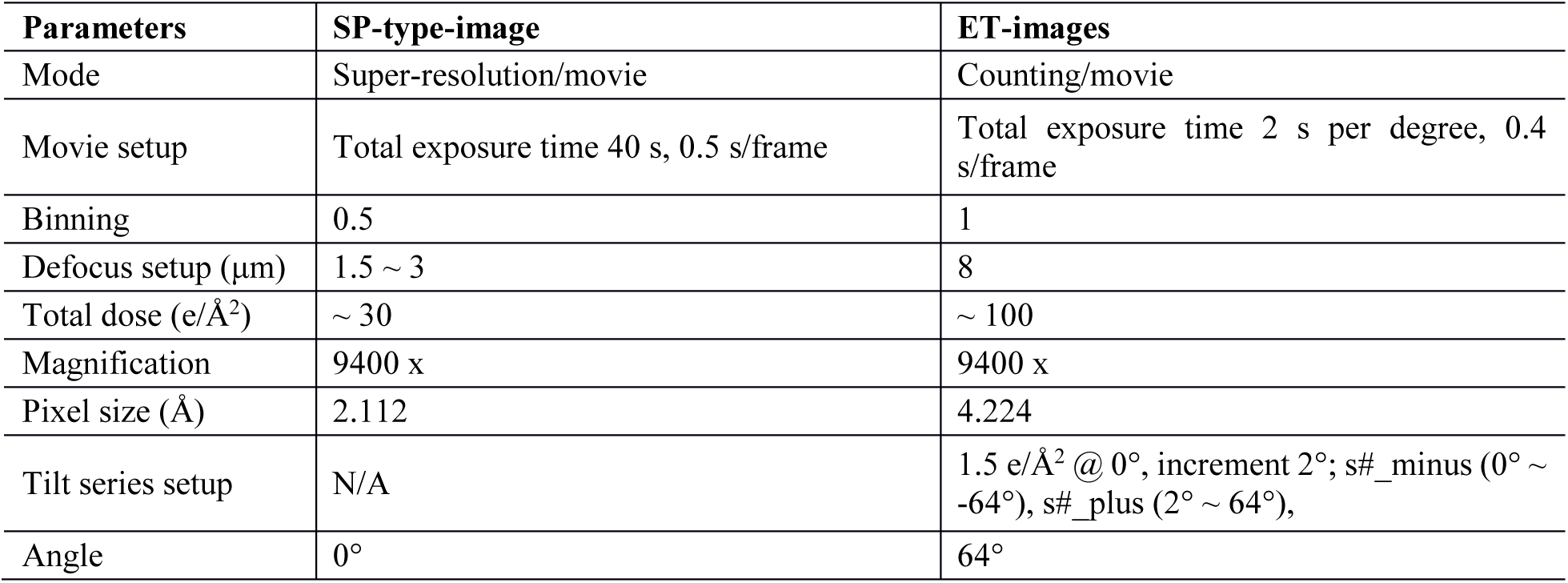
K2 parameters setup for TYGRESS.

| Parameters | SP-type-image | ET-images |
| --- | --- | --- |
| Mode | Super-resolution/movie | Counting/movie |
| Movie setup | Total exposure time 40 s, 0.5 s/frame | Total exposure time 2 s per degree, 0.4 s/frame |
| Binning | 0.5 | 1 |
| Defocus setup (μm) | 1.5 ~ 3 | 8 |
| Total dose (e/Å<sup>2</sup>) | ~ 30 | ~ 100 |
| Magnification | 9400 x | 9400 x |
| Pixel size (Å) | 2.112 | 4.224 |
| Tilt series setup | N/A | 1.5 e/Å<sup>2</sup> @ 0°, increment 2°; s#_minus (0° ~ -64°), s#_plus (2° ~ 64°), |
| Angle | 0° | 64° |

